# Migrating in a warming world: A deep learning approach to predict pan-American seasonal shifts in the monarch butterfly niche

**DOI:** 10.1101/2025.06.25.661455

**Authors:** Chiara Vanalli, Robin Zbinden, Nina van Tiel, Devis Tuia

## Abstract

Climate change is driving biodiversity loss, disrupting ecosystem functioning, and altering species distributions. Migratory species, whose range varies across seasons depending on specific climatic conditions, are particularly sensitive to environmental changes and serve as indicators of ecosystem health. However, current species distribution models often fail to capture the temporal dynamics critical for migratory species, limiting their ability to provide accurate future range estimations. In this study, we address this gap by developing a time-aware deep learning species distribution model for the monarch butterfly (*Danaus plexippus*), an iconic species for biodiversity conservation. Using monarch occurrence records across the Americas gathered from scientific and citizen science sources, we embed the effect of monthly climatic variables in a sequential framework. We compare the performance of our seasonal model to conventional time-static baselines, showing not only better performance in the present, where the models have been trained and validated, but also in the past. Our findings show that climatic factors such as humidity, temperature, precipitation and cloud coverage strongly influence the ecological niche of the monarch butterfly, with notable seasonal and spatial variability. Applying our model under climate change scenarios, we predict a northwestward shift in the monarch range by the end of the XXI century, with expansion in Canada and significant contraction in California and Central America, key sites for overwintering that also host resident monarch populations. These changes could severely impact the species migratory cycle and population stability. Using Shapley values, an explainable AI technique, we identify the decrease in precipitation and humidity as important environmental drivers responsible for the contraction of overwintering sites. By focusing on a species of high ecological relevance through a time-aware modeling approach, this work brings novel insights for the conservation of migratory species in the face of the challenges posed by climate change.

## Introduction

Climate change is profoundly affecting ecosystem health by altering weather patterns and frequency of extreme events, with disruptive consequences on the ability of species to survive, reproduce, and thrive [1, 2]. The ongoing pervasive decline in biodiversity serves as a clear signal of the threat that anthropogenic pressures pose to ecosystems [3]. The degradation of ecosystems and the essential services that they provide undermines human well-being with significant economic and societal costs [4, 5, 6]. These challenges highlight the urgent need to deepen our understanding of the link between species and the environment in which they live, as well as to develop innovative tools that can track and predict the evolving status of ecosystems in response to environmental changes. By translating these insights into actionable strategies, science-informed conservation measures can effectively enhance the resilience of ecosystems and safeguard the diversity of the species they host.

Migratory species play a key role in ecological networks, by transporting nutrients and impacting trophic webs, therefore providing important ecological services along their migratory route [7, 8]. Whether driven by survival or reproduction, migrations are an adaptive response to specific environmental conditions and, in turn, highly impacted by environmental changes [9, 8, 10]. Global warming has contributed to earlier migration and breeding time [11], shorter migration distances [12], and loss of breeding ground [13] with detrimental alterations to the migration journey. Therefore, understanding how migratory species distributions are shaped by climatic conditions and how they are expected to shift due to climate change is key for ecosystem and biodiversity conservation [7]. In this respect, ecological niche modeling can provide crucial insights, identifying the range occupied by a species in relation to environmental variables [14, 15] and its expansion, contraction, and shift due to climate change [16, 17]. However, mapping the distribution of migratory species to the environment is challenging due to the intrinsic spatial and temporal connectivity that characterizes migration: gaps in tracking individual movements throughout the entire migratory cycle and the overlook of the complex impacts of the multiple seasonal climatic factors represent the main limitations and lead to the current partial success of modeling migratory patterns in space and time at high resolution.

Recent technological advancements have brought new opportunities for species monitoring, such as the use of positional transmitters (GPS) [18, 19, 20] and drones with cameras [21, 22]. Concurrently, the growing amount of georeferenced species observations collected from the general public worldwide and shared through citizen-science platforms represents a powerful resource for ecological and environmental sciences [23], but also for tackling planetary health challenges [24]. Leveraging these novel data streams, Species Distribution Models (SDMs) have become a widely used approach to map species niches by statistically linking observed species presences/absences to environmental conditions [25, 26, 27]. Despite their impressive achievements in ecological niche mapping, SDM applications to migratory species are limited [28, 29]. This is mainly due to the time-static way in which environmental inputs (usually averaged over multiple decades) and species occurrence probability are treated, neglecting the temporal dynamics that are pivotal for migratory species. The diverse climatic conditions to which these species are exposed at different stages of their migratory route need to be thoroughly addressed and embedded in modeling frameworks [30, 31]. Disregarding those seasonal dynamics could result in an over- or under-estimation of the actual species distribution [30], hindering the model implementation under different climatic conditions from those where the model has been trained and validated [32] thus limiting their ability to draw meaningful conclusions on species ecology [33]. Deep learning-based SDMs (deep SDMs) have the potential to overcome the highlighted limitations, thanks to their flexibility to treat large and diverse input information [34, 35], handle complex nonlinear relations, proving to be robust and effective tools to model species distribution [36, 37, 38, 39, 40] and more generally to inform biodiversity conservation [41, 42]. Additionally, advancements in explainable Artificial Intelligence methods [43, 44] have made it possible to transition black box models into approaches capable of identifying the relevant predictors reflecting the ecological dynamics of the target species and/or explaining observed shifts [34, 45]. Therefore, incorporating the temporal dimension has the potential to *i*) improve their biological realism of deep SDMs, properly addressing the different role of predictors at different time periods, *ii*) reduce biases in niche estimates, possibly due to the mismatch of specific environmental predictors, *iii*) produce a time-dynamic species occurrence probability that is key to follow the journey of migratory species, and *iv*) enhance model robustness to predict species range alterations under rapid global change, as informed by general circulation models.

In this work, we aim to map the ecological niche of the monarch butterfly (*Danaus plexippus*) at a pan-American scale and to predict future projections of the niche shifts under climate change. A key pollinator species [46] with a migratory route that extends up to 4,000 km, monarchs have historically been an iconic migratory species for biodiversity conservation [47, 48] and, together with other butterfly species, can be used as sentinel to monitor ecosystem health [49, 50]. Climatic factors, habitat loss, diseases, and agricultural insecticide use have threatened monarch populations in the Americas, with sharp declines over the last three decades [51, 52]. In particular, climatic changes have been identified as a major driver of monarch population dynamics with significant impacts on the breeding-season range and population size [13]. In 2023, the International Union for Conservation of Nature modified the conservation status of the monarch butterfly from ‘endangered’ to ‘vulnerable’ to extinction, after detecting a stable/positive population trend [53], despite some criticisms from the scientific community, who showed that evidence of growing monarch population was not statistically significant [54]. In this study, we used large-scale occurrence observations of monarchs, collected from both professionals and the general public, to examine the role of detailed (monthly) climatic inputs for predicting their seasonal distribution across the Americas. We developed a time-aware deep SDM that captures the temporal dynamics as seasonal sequences, and compared its performance to a baseline time-static model. For each season, we assessed the different contributions of the multiple climatic variables to the definition of species ranges, using an explainable AI approach. Last, we predicted future contractions/expansions and shifts of the monarch ecological niche under different climate change scenarios, including the uncertainty generated by the multiple scenarios considered.

Evaluating how climate change shapes the ecological niche of a well-studied species, like the monarch butterfly, with a complex ecology and migration, provides valuable insights into macroecology and biogeography of threatened species. Our approach shows clear predictive benefits, leading to a better outlining of the niche occupied by the Monarch butterfly. Our predictions for the present status and possible future ranges hold the potential of better informing science-based conservation efforts for habitat protection and/or restoration to face the challenges posed by global warming.

## Material and Methods

### Study system: Monarch butterfly in the Americas

The monarch butterfly (*Danaus plexippus*) is native to North and South America, where it exhibits a wide diversity of migratory behaviors [55]. The two main long-distance migratory monarch populations are: the North Eastern American population, which every year travels 4,000 km in a multi-generational manner to overwinter in mountainous forests in Mexico [56], and the North Western American population, which is smaller in size and also migrates in the fall, traveling up to hundreds of km to forested groves along the coast of California [57, 58]. This migratory behavior is driven by diapause, a hormonally controlled suspended reproductive state that contributes to winter survival and to the recolonization of northern territories [59, 60]. In addition to these migratory populations, non-migratory stable populations of monarchs have been documented in southern California [61, 62], southern Florida [63], the Gulf Coast [64] and Central America [65]. Seasonal climate variability is the main driver of monarch migration [66, 60] and recent changes in climate related to global warming have been impacting monarch butterfly dynamics and distribution [13]. The diversity of the monarch migratory behaviors, with seasonal and year-round suitable habitat across the Americas makes studying and modeling climate change impacts challenging. In addition to the Americas, *D. plexippus* also inhabits different areas of the world (i.e. New Zeland, Australia and some islands of south western Europe [55]), but for the purpose of this study we focus on the main populations in the Americas within the spatial domain that spans from -10° to 65° of latitude and from -130° to -38° of longitude.

### Datasets: Species occurrence and climatic predictors

We used presence-only data of *D. plexippus* with georeferenced coordinates available on the Global Biodiversity Information Facility (GBIF) [67] from 2010 to 2024 to train, validate, test, and compare performance of the models described (data random split of 80% for training, 10% for validation, and 10% for testing). We considered observations recorded in *i*) March, April, and May belong to the spring season, *ii*) June, July, and August to the summer season, *iii*) September, October, and November to the fall season, and *iv*) December, January, and February to the winter season, following the seasonal labeling of the boreal hemisphere. The choice of the selected 15-year time span is justified by the temporal coverage of observations with available seasonal information (Figure A1), as well as an appropriate time scale to examine the effects of climatic variables on the ecological niche of monarchs. In addition, we use *D. plexippus* presence-only data from 1990 to 2004 that lack seasonal information (i.e. the observations do not include the temporal detail about the month of observation, see Figure A1). Given the lack of temporal information, we validate our simulations for the period 1990-2004 only at the yearly level and analyze the seasonal trends only qualitatively.

The abiotic variables used in our deepSDM are: minimum (*tasmin*), mean (*tasmean*) and maximum (*tasmax*) temperature, specific (*huss*) and relative humidity (*hurs*), evaporation (*evapsbl*), downwelling shortwave (*rsds*) and longwave radiation (*rlds*), wind speed (*sfcwind*), precipitation (*pr*), cloud coverage (*clt*) (Table A1). They are available from the Coupled Model Intercomparison Project Phase 6 (CNRM-CM6-1-HR model) at a spatial resolution of ∼ (0.5 × 0.5) degrees, for the past, present and future climate under different Shared Socioeconomic Pathways (SSPs) of the Intergovernmental Panel of Climate Change [68]. Climatic data were downloaded at a monthly climatic resolution and averaged over the considered 15-year temporal horizon, therefore providing data for each month averaged over the entire time period. Specifically, we analyze a more sustainable scenario, SSP1-2.6, an intermediate emission scenario, SSP2-4.5, and a fossil-fuel oriented scenario, SSP5-8.5, over a 15-year time period at the middle of the century, from 2046 to 2060 (Figure A2), and at the end of the century, from 2086 to 2100 (results in the main manuscript).

### Seasonal Species Distribution Model

Extensive modeling efforts have been made towards the understanding of the *D. plexippus* ecological niche [69, 66, 70] as well as towards the prediction of its future range shifts [71, 72, 73]. Despite the recognized importance of incorporating seasonality [73, 30], the developed SDMs for the monarch butterfly often focus on a restricted spatial domain [73, 74, 75] and/or on a specific time interval of the yearly migration cycle [69, 70] and/or examine only few climatic predictors, usually temperature and precipitation [75, 73]. Conversely, a more holistic framework that includes dependence upon multiple climatic variables, across the entire spatial domain of its migratory route may enable a better understanding of climate change impacts upon the monarch butterfly seasonal niche and the differences across stable and migratory populations. Therefore, we incorporate the 11 climatic predictors (Table A1) across the pan-American continental scale as inputs to a series of Multi-Layer Perceptrons, artificial neural networks previously implemented in species distribution modeling [38, 76]. Specifically, we compare three different modeling settings (Figure 1): *i*) **a time-static model (M0)** that takes as input annual average predictors across the study period (total of 11 predictors) and produces the probability of monarch occurrence in a time-independent manner, *ii*) **seasonal independent models (M1)** that incorporate monthly climatic predictors within the considered season (each season being composed of three months, this leads to a total of 3×11 predictors for each season) and produce the seasonal probability of monarch occurrence and *iii*) **seasonal concatenated models (M2)** that considers as inputs, in addition to the monthly climatic predictors, the estimated probability of occurrence of the previous season, therefore propagating niche information across seasons. The seasonal models can be concatenated and run sequentially. We constrained this chain of models to the three previous seasons (e.g. winter probability is estimated starting from the previous spring season). For the first season in the concatenation (spring in the previous example), the previous season probability is set to 0.5 in all locations. Finally, we input an extra dummy starting input value set to 1 if the considered season is the start of the concatenation or to 0, otherwise. Overall, this setting corresponds to a total of 3×11+2=35 predictors for each season, where the two extra variables are the occurrence probability predicted for the preceding season and the dummy variable, respectively. The same architecture is kept across the four seasons, with two hidden layers of 40 neurons each and one output node, corresponding to the *D. plexippus* occurrence probability for the season considered. As nonlinearities, ReLU activation functions are implemented throughout the architecture, and a final sigmoidal transformation is employed to predict probability values between 0 and 1. We generate 10,000 background points [77] in a buffer of 100 km around the convex hull defined by species observations. These are used together with presence data to train, validate, and test the M0, M1, and M2 models. Cross-entropy loss for binary classification and Adam optimizer [78], with a learning rate of 0.01 and 250 epochs, are used for model training. We assess the model performance on the test set computing the area under the ROC curve (*AUC_ROC_*), the area under the precision-recall curve (*AUC_P R_*), and the true skill statistic (*TSS*). Since the latter is a threshold-dependent metric that requires binary model outputs, we used the validation dataset to find the optimal thresholds that maximize *TSS*. During the time period 2010-2024, the global (i.e. any season) model performance is computed by averaging the four seasonal performances. In this same time period, we examined the performance of the Maximum Entropy method (MaxEnt), a common machine learning approach to model species distribution, using linear, hinge and product transformations of time-static features [79]. Instead, given that no seasonal information is available for past observations (1990-2004), only a time-static performance is evaluated. In this period, to compare occurrence probabilities generated with different seasonal models with contrasting binarization thresholds for the global *TSS*, we rescale the seasonal probabilities according to a 0.5 threshold: *p_rescaled,i_* = *p_i_* − (*th_i_* − 0.5), where *th_i_* is the season-specific probability threshold for binarization.

**Figure 1:**
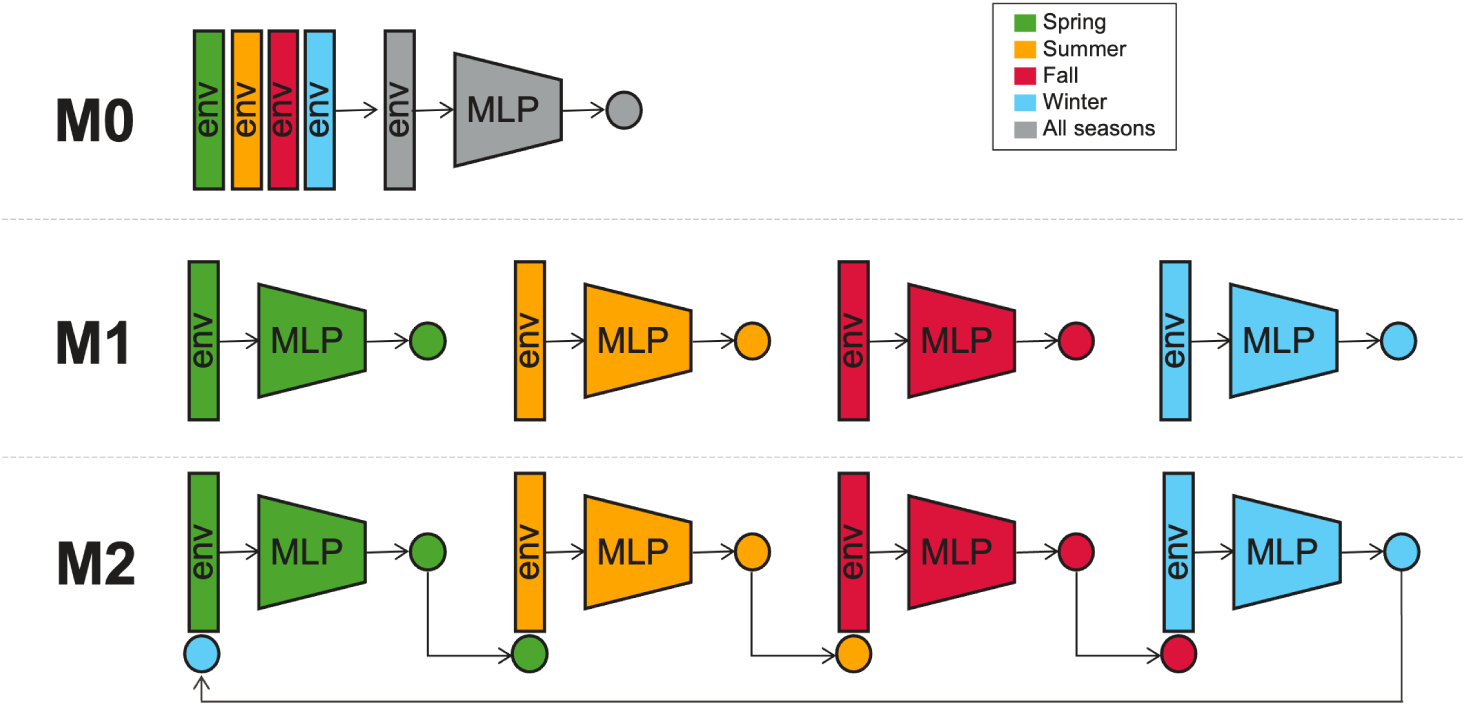
Time-static (M0), seasonal independent (M1), and seasonal concatenated (M2) analyzed species distribution models. Monthly environmental predictors (env, Table A1) are used as inputs to Multi Layer Perceptrons (MLP) with monarch occurrence probability as output node. Spring season is represented in green, summer in orange, fall in red and winter in cyan, while M0 is represented in gray because it takes as input annual average predictors and produces the probability of monarch occurrence in a time-independent manner.

To examine the contribution of each climatic predictor in defining the monarch butterfly niche, we consider the Shapley values [80], an eXplainable Artificial Intelligence (XAI) method [44, 34]. For each considered feature and analyzed season, we calculate Shapley values, offering a measure of variable importance to the species occurrence, together with the Pearson correlation between Shapley and feature values, whose sign and magnitude represents the overall directionality effect of the focal feature on the species probability of occurrence [45]. Despite providing general information on the feature effects, the latter method is a simplification which assumes linear relationships between feature values and Shapley values, although this is not necessarily the case. Last, we assess the difference in Shapley values between the future scenarios and the present situation to examine how changes in specific climatic predictors impact the occurrence of the monarch butterfly. While we calculate Shapley values for all the considered environmental predictors, we explore the spatial distribution of selected well-studied variables, such as humidity, precipitation and temperature for specific seasons.

## Results

### Model comparison and data fitting

We compare the performance of the time-static model (M0), seasonal independent (M1), and concatenated models (M2) in the present for which we have seasonal labels (2010-2024, Tables 1 and A2) and in the past for which no seasonal labels were available (1990-2004, Table 2). The time-static baseline already shows good global performance, with an AUC of 0.874 and a TSS of 0.683, confirming the important role that climatic predictors play in shaping the distribution of monarch butterflies [13]. A comparable model performance is achieved by MaxEnt with time-static environmental inputs, suggesting that both analyzed approaches (i.e., MLPs and MaxEnt) are able to represent well the environmental niche of the monarch butterfly (Table A2). Model performance achieved with MLPs is even more evident for seasons with higher sample size (Figure A1), such as summer and fall, for which M0 achieves AUC values higher than 0.90. However, M0 shows a weaker performance in accurately modeling the winter distribution, a key task to identify potential overwintering sites that are essential to population preservation and migration. Deploying independent seasonal models (M1) partially solves this issue, with a greater increase in model performance in winter (+14%) than in summer (+1%) compared to M0. This gain in model accuracy, given by the integration of monthly climatic predictors, is an indicator of the importance of seasonality for migratory species like the monarch butterfly. Compared to M1, the seasonal concatenated model (M2) presents just one more input (i.e. the probability of occurrence of the previous season), together with a dummy variable, but still results in an increase in the model performance. This increase is consistent across seasons, but more pronounced for spring and summer seasons and depends on the estimated occurrence probability of winter and spring, respectively. When these same models are validated in the distant past, although we do not have seasonal labels, similar positive trends in model performance are found in particular for TSS and AUC_ROC_, which experience a gain of +15% and +2.5%, respectively, when comparing M2 to M0 (Table 2). M2 achieves the best model performance and more realistically represents the link between different seasonal stages of the migratory cycle, which is confirmed by its ability to better reproduce the niche in the distant past. Therefore, we select the concatenated seasonal model architecture (M2) to project future shifts of the niche of *D. plexippus* under climate change.

**Table 1:** Global (any season) and seasonal model performance comparison of the Area Under the ROC Curve (AUC_ROC_), the Area Under the Precision-Recall Curve (AUC_PR_), and the True Skill Statistic (TSS) using the test dataset of 2010-2024. M0 represents the time-static model, M1 the seasonal independent model, and M2 the seasonal concatenated model, respectively. The percentage increase in model performance compared to the time-static model M0 is reported in square brackets.

| | Model | $AUC_{ROC}$ | $AUC_{PR}$ | $TSS$ |
| --- | --- | --- | --- | --- |
| Global | M0 | 0.874 | 0.903 | 0.683 |
|  | M1 | 0.927 [+6.1%] | 0.974 [+7.9%] | 0.781 [+14%] |
|  | M2 | 0.944 [+8.0%] | 0.978 [+8.3%] | 0.806 [+18%] |
| Spring | M0 | 0.862 | 0.893 | 0.688 |
|  | M1 | 0.921 [+6.8%] | 0.967 [+8.3%] | 0.777 [+13%] |
|  | M2 | 0.944 [+9.5%] | 0.973 [+9.0%] | 0.813 [+18%] |
| Summer | M0 | 0.907 | 0.980 | 0.687 |
|  | M1 | 0.916 [+1.0%] | 0.988 [+0.8%] | 0.715 [+4.1%] |
|  | M2 | 0.943 [+4.0%] | 0.991 [+1.1%] | 0.770 [+12%] |
| Fall | M0 | 0.886 | 0.966 | 0.684 |
|  | M1 | 0.912 [+2.9%] | 0.978 [+1.2%] | 0.763 [+12%] |
|  | M2 | 0.925 [+4.4%] | 0.982 [+1.7%] | 0.771 [+13%] |
| Winter | M0 | 0.840 | 0.771 | 0.674 |
|  | M1 | 0.960 [+14%] | 0.962 [+25%] | 0.869 [+29%] |
|  | M2 | 0.963 [+15%] | 0.964 [+25%] | 0.869 [+29%] |

**Table 2:** Global (any season) model performance comparison of the Area Under the ROC Curve (AUC_ROC_), the Area Under the Precision-Recall Curve (AUC_PR_) and the True Skill Statistic (TSS) using the time-static dataset of 1990-2004. M0 represents the time-static model, M1 the seasonal independent model, and M2 is the seasonal concatenated model, respectively. Percentage increase in model performance compared to the time-static model M0 is reported in square brackets.

| | Model | $AUC_{ROC}$ | $AUC_{PR}$ | $TSS$ |
| --- | --- | --- | --- | --- |
| Global | M0 | 0.846 | 0.989 | 0.619 |
|  | M1 | 0.864 [+2.1%] | 0.989 [+0%] | 0.658 [+6.3%] |
|  | M2 | 0.867 [+2.5%] | 0.990 [+0.1%] | 0.713 [+15%] |

Mapping the seasonal distribution of monarchs at a pan-American scale, we find a good overlap between the estimated niche and the occurrence data (Figure 2). Importantly, the seasonal niches represent well the monarch migratory route, with the summer distribution extending the most northwards and the most concentrated niche in overwintering sites. Our model is able to identify suitable areas that allow monarch presence year-round, which match well with known areas of stable monarch populations, such as southern California [61, 62], southern Florida [63], the Gulf Coast [64], and Central America [65] (Figure 2b,d). Our findings can indeed clarify which climatic conditions favor the stability versus the migratory behavior of monarch butterfly populations. However, our model performance could be improved in the Midwest of the United States, which does not belong to the estimated niche despite some observations having been recorded. Ultimately, our time-aware modeling approach reproduces well the observed occurrences of monarchs and can be effectively used for projecting the ecological niche in different temporal and climatic settings.

**Figure 2:**
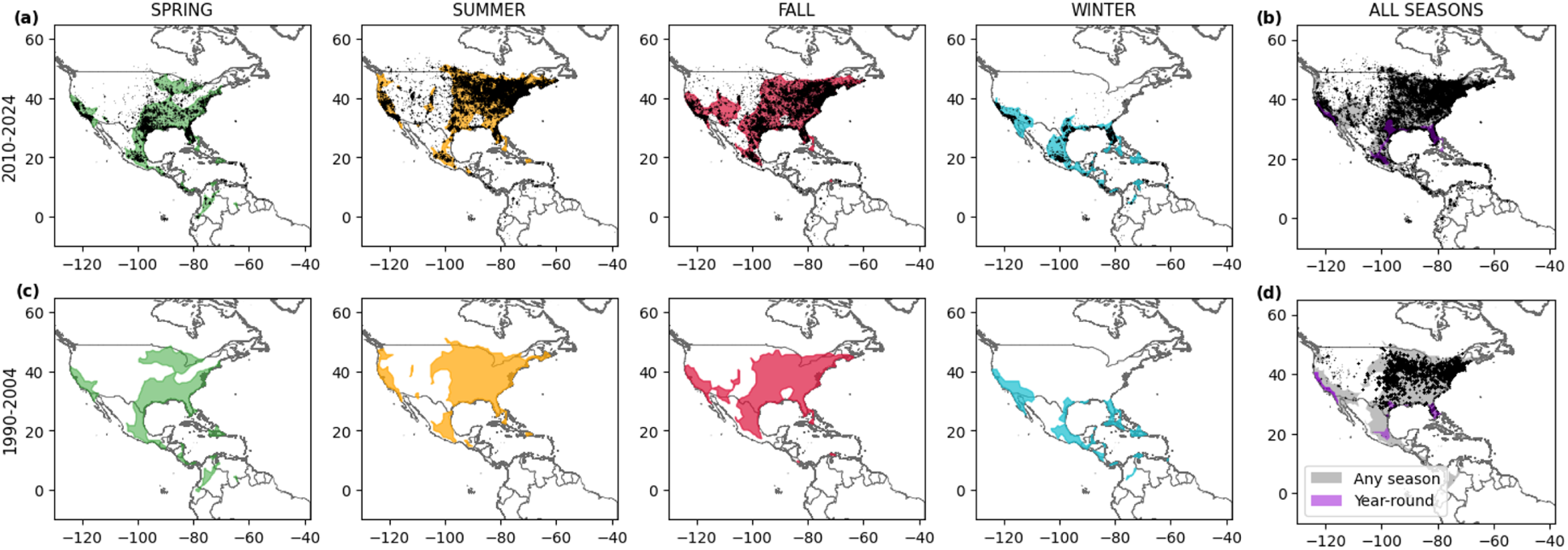
Estimated seasonal (a, c) and global (all seasons, b, d) niches of the monarch butterfly (spring in green, summer in orange, fall in red, winter in cyan, any season in gray, and year-round in purple) by the concatenated seasonal model (M2) with monarch observations in black in the present (2010-2024, a, b) and past (1990-2004, c, d). Note that the observation records for the time period 1990-2004 lack seasonal information, but we can produce seasonal maps thanks to the M2 model architecture.

### Contribution of climatic predictors

We quantify overall predictor importance using the mean of absolute values of Shapley values (Figure 3a) and estimate the directional effect of the predictors on the species probability of occurrence using Pearson correlation coefficients (Figure 3b). Shapley values allow us to interpret model predictions by examining the concurrent contributions of climatic features to species occurrence in a spatiotemporally-explicit way. Comparing the importance of each seasonal climatic predictor, relative (*hurs*) and specific humidity (*huss*), and downwelling longwave radiation (*rlds*) seems to be highly positive correlated with Shapley values. Interestingly, the predictor importance varies across seasons, which may result in an inversion of the sign of the correlation coefficient. Overall, cloud coverage and downwelling longwave radiation are consistently negatively associated with the monarch butterfly occurrence, whereas precipitation and relative humidity are positively related to species presence. The probability of occurrence of the previous seasons is positively correlated with monarch presence, in particular in spring and summer, and weakly negatively correlated in winter, possibly due to the uniqueness of the winter distribution. The correlation is weaker in fall, probably due to the drastic shift occurring when reaching the overwintering sites. Temperature-related predictors (*tas*, *tasmin* and *tasmax*) positively affect monarch occurrences in winter. These results highlight the complex and multi-faceted impact of climatic predictors in defining the monarch butterfly occurrence, which becomes even more heterogeneous when we analyze the effect in a spatially explicit way of selected key predictors (Figure 3c,d). In particular, we focus on the Shapley values of average temperature (*tas*) in summer and winter, precipitation in spring (*pr*) and relative humidity in summer (*hurs*). The importance of drastic temperature shifts from summer to winter is evident, leading to high Shapley values located mostly in Central America, where temperatures are above 15 *^◦^*C. Shapley values of spring precipitation are high in the East and West coasts of the US, where precipitation is more abundant, compared to the drier central regions. The same pattern, but more extreme, is outlined by humidity in summer. The different contribution of climatic predictors across seasons and spatial locations highlights the need for season-aware species distribution models that can be flexibly implemented for the multiple temporal stages of the migration cycle.

**Figure 3:**
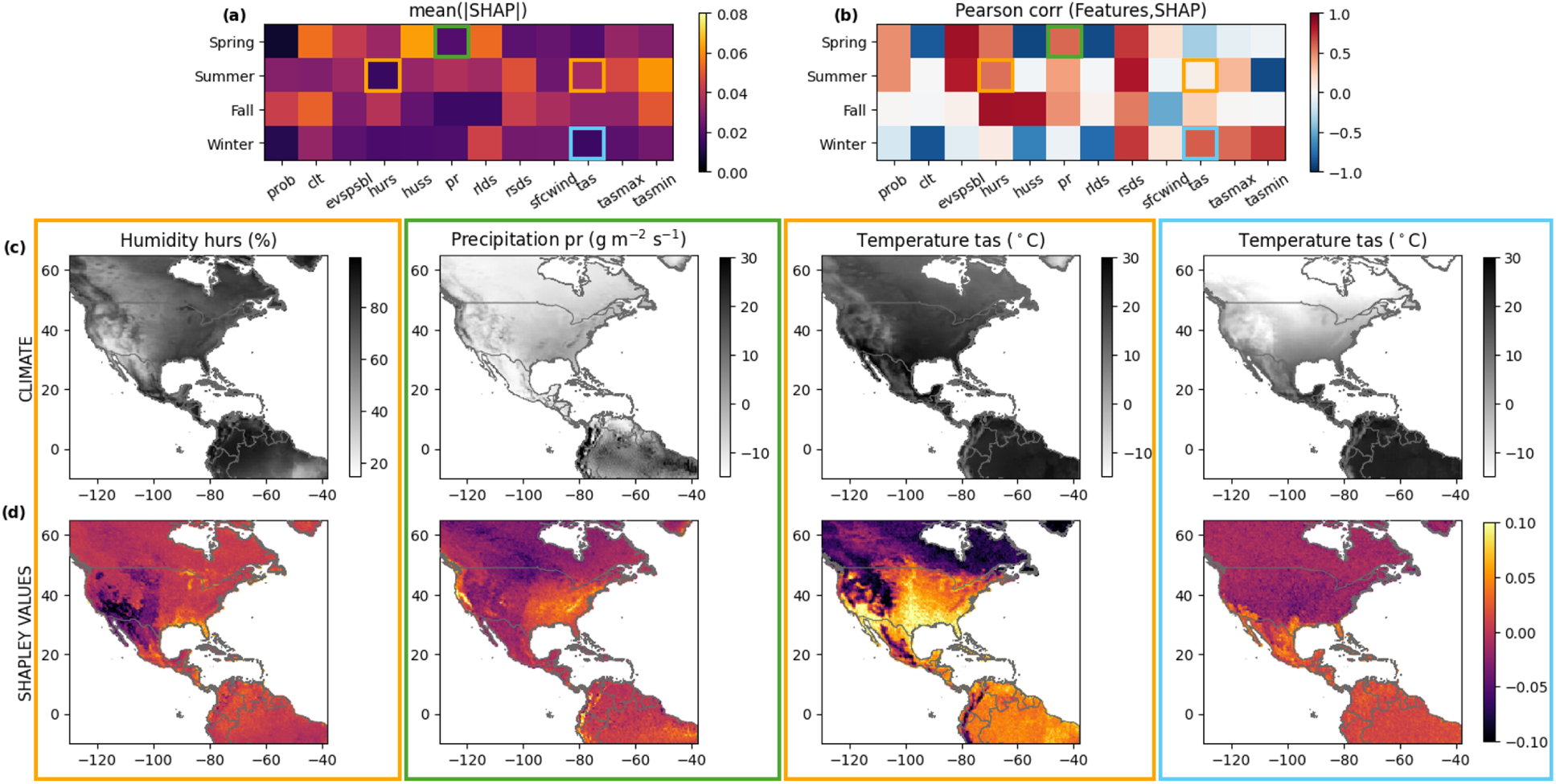
(a) Seasonal mean absolute Shapley values for each climatic predictor and (b) Pearson correlation coefficients between features and Shapley values. Spatial maps of (c) climatic variables and (d) Shapley values of relative humidity in summer (*hurs*, %), precipitation in spring (*pr*, g m^-2^ s^-1^), and temperature in summer and winter (*tas*, *^◦^*C). The colored boxes highlight the climatic variables that are investigated. Their color represents the considered season: spring (green), summer (orange), fall (red), winter (cyan). Refer to Table A1 for variable abbreviations and full description.

## Future shifts in the monarch ecological niche

We project future seasonal and season-independent niches of the monarch butterfly at the middle (Figure A2) and at the end of the XXI century (Figure 4) under multiple climate change scenarios. Going from the mildest (SSP1-2.6) to the most extreme (SSP5-8.5) scenario, we notice more evident distributional shifts. Our model predicts a significant gain in new suitable areas, which is more significant in summer and fall. These new suitable areas would be mainly located on the East Coast for spring, in the northern part of the US and Canada for summer and fall, and in the southern US for winter. This shift would contract the spring-summer niche areas in Central America. Overwintering sites would also expand northwards, covering up to the entire southern US. These expected changes in the monarch seasonal distributions can potentially have significant consequences on the suitable areas for year-round butterfly populations. In this respect, our model predicts the disappearance of the stable populations in Central America and California, in particular, under the SSP1-2.6 (Figure 4a) and SSP5-8.5 (Figure 4e) scenarios. This important loss is also predicted for the middle of the century (Figure A2), depicting an alarming situation given that these populations can fail to adapt their resident behavior to a migratory one. The greatest losses would be experienced under the SSP5-8.5 pathway and are most marked in spring and for resident populations, and would be located in Central America, California, and Florida. Even though with a smaller extension, these areas are indicated to become unsuitable even under milder climate change scenarios. Conversely, an expansion of the resident monarch population in the southern states of the US, such as Texas, Georgia, Louisiana, and Alabama, is projected (Figure 4g). According to the simulations, these regions may represent the only areas where it would be possible to find monarchs year-round in the future.

**Figure 4:**
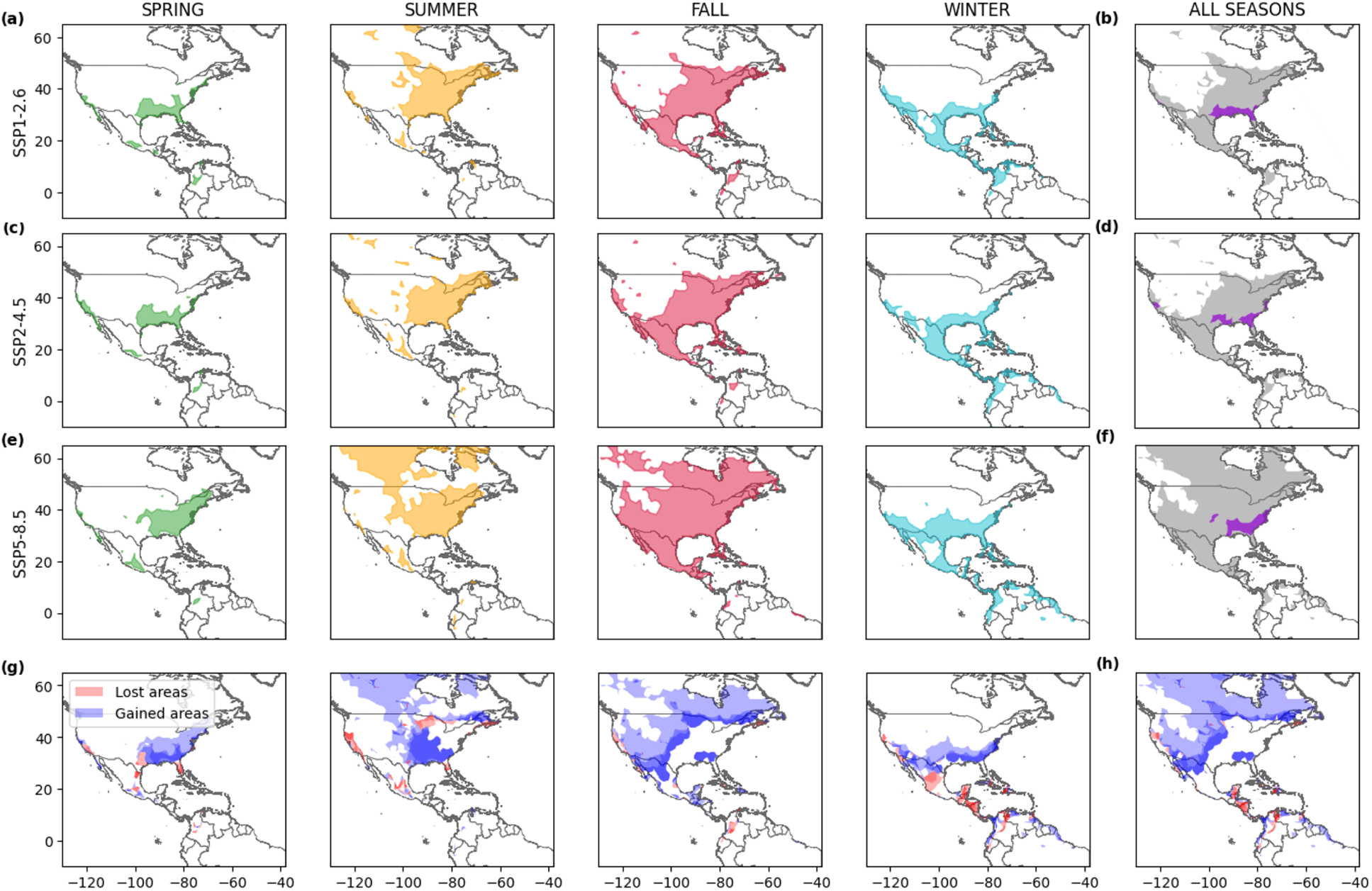
Projected seasonal seasonal (a, c, e) and global (all seasons, b, d, f) niches of the monarch butterfly (spring in green, summer in orange, fall in red, winter in cyan, any season in gray and year-round in purple) at the end of the XXI century (2086-2100) under SSP1-2.6 (a, b), SSP2-4.5 (c, d) and SSP5-8.5 (e, f) climate change scenarios. Lost (red) and gained (blue) areas, compared to 2010-2024, are represented for each season (g) and for all seasons (h). Darker areas (panels g, h) represent regions of agreement between the three scenarios.

Analyzing the latitudinal and longitudinal shifts of the monarch ecological niche (Figure 5a,b and Figure A3), we expect a median northward shift of +0.5 or +9 degrees and a western shift of -1.5 or -6 degrees for SSP1-2.6 or SSP5-8.5, respectively. The most prominent latitudinal shifts are predicted for the upper limits of the summer and fall niches, with +17 and +12 degrees of the 95^th^ percentile, respectively (Figure A3). Longitudinal eastern shifts of the spring distribution, due to the contraction and loss of suitable areas on the West Coast, would result in the same directional shift of the year-round niche. The highlighted climate-driven shifts would result in the loss of suitable areas for monarchs, ranging from 10% to 85% of the current niche, compared to the present (Figure 5c). Although the estimated extension of niche expansion areas outnumbers the lost ones (Figure 5d), the ability of monarchs to reach, adapt, and thrive in those new areas remains unknown.

**Figure 5:**
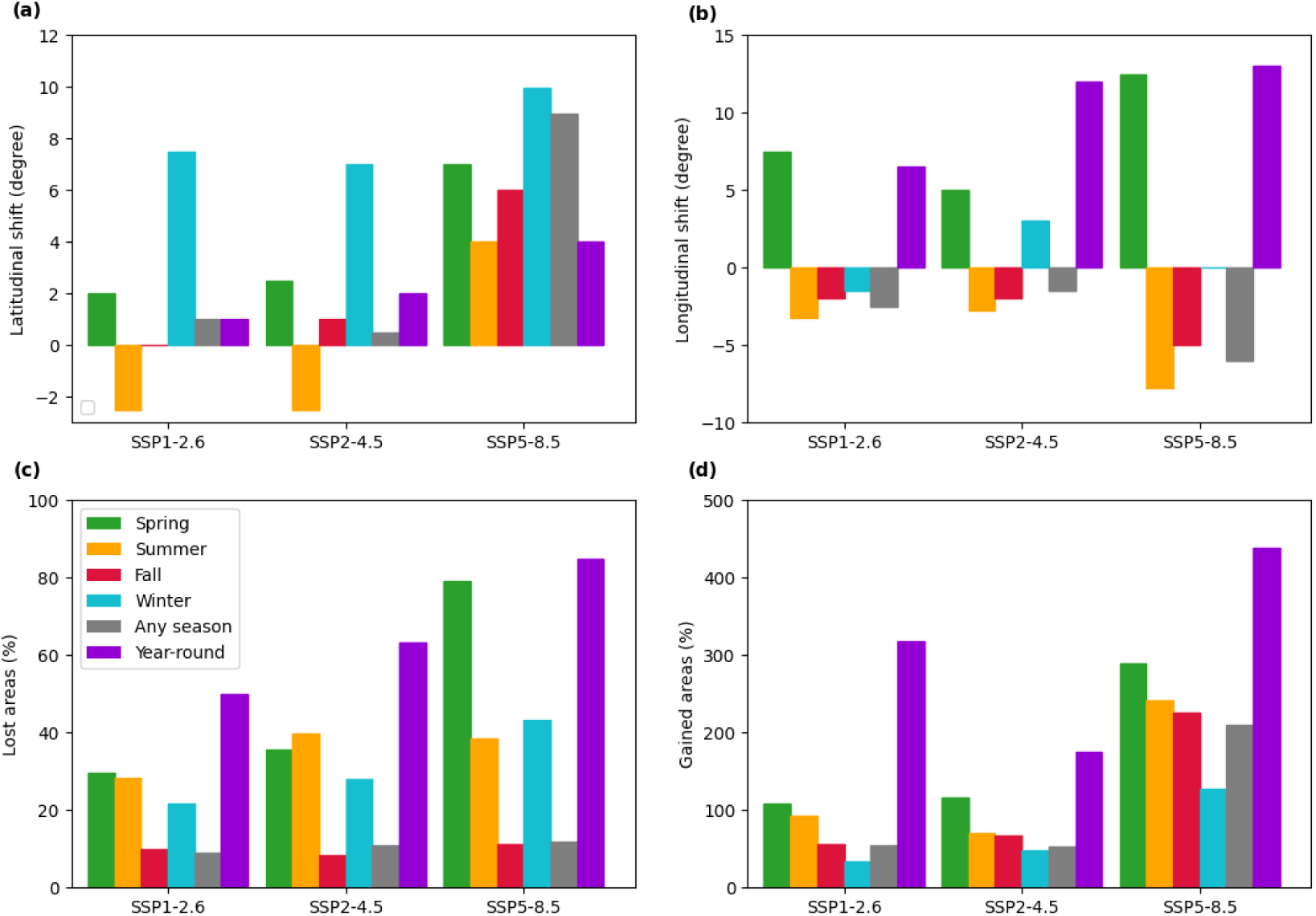
(a) Latitudinal and (b) longitudinal future shifts of the medians of the monarch spatial distribution at the end of the XXI century (2086-2100) under SSP1-2.6, SSP2-4.5 and SSP5-8.5 climate change scenarios compared to the present (2010-2024) for spring (green), summer (orange), fall (red), winter (cyan), any season (gray) and for year-round (purple). Percentages of (c) lost and (d) gained areas of the monarch butterfly ecological niche are reported. Refer to Figure A4 for detailed information about latitudinal and longitudinal spatial distribution.

Thanks to the implementation of the concatenated seasonal deep SDM modeling approach, not only can we visualize the expected future niche shifts described above, but also determine which shifts in climatic drivers are responsible for producing these patterns. Computing the average difference between Shapley values in the future compared to the selected historical baseline (2010-2024), it becomes possible to disentangle the role of the multiple model predictors in determining the niche shifts (Figure 6). At the end of the century, under the SSP5-8.5 scenario, a positive difference in Shapley values is found for temperature predictors in winter, indicating a higher suitability for monarchs, probably due to less restrictive thermal conditions. Evapotranspiration is expected to be less favorable for monarch occurrence, in particular in the warmer seasons of the year (Figure 6b). Less prominent, but similar trends across the three considered scenarios are expected at the middle of the century (Figure A4). Exploring changes in Shapley values spatially (Figure 6c,d), we find an increase in Shapley values in the north-central US for summer temperature, contrasted by a decrease in Central America. In contrast, an opposite increase in Shapley values is highlighted for the central territories for winter temperature. For precipitation, a consistent decrease in Shapley values is expected in California and the Gulf Coast, due to drier conditions. These latter changes, together with the combination of the other climatic predictors, are responsible for the future contraction of the year-round niche of monarch butterflies. The described trends are expected to be milder under SSP1-2.6 and SSP2-4.5 (Figures A5, A6).

**Figure 6:**
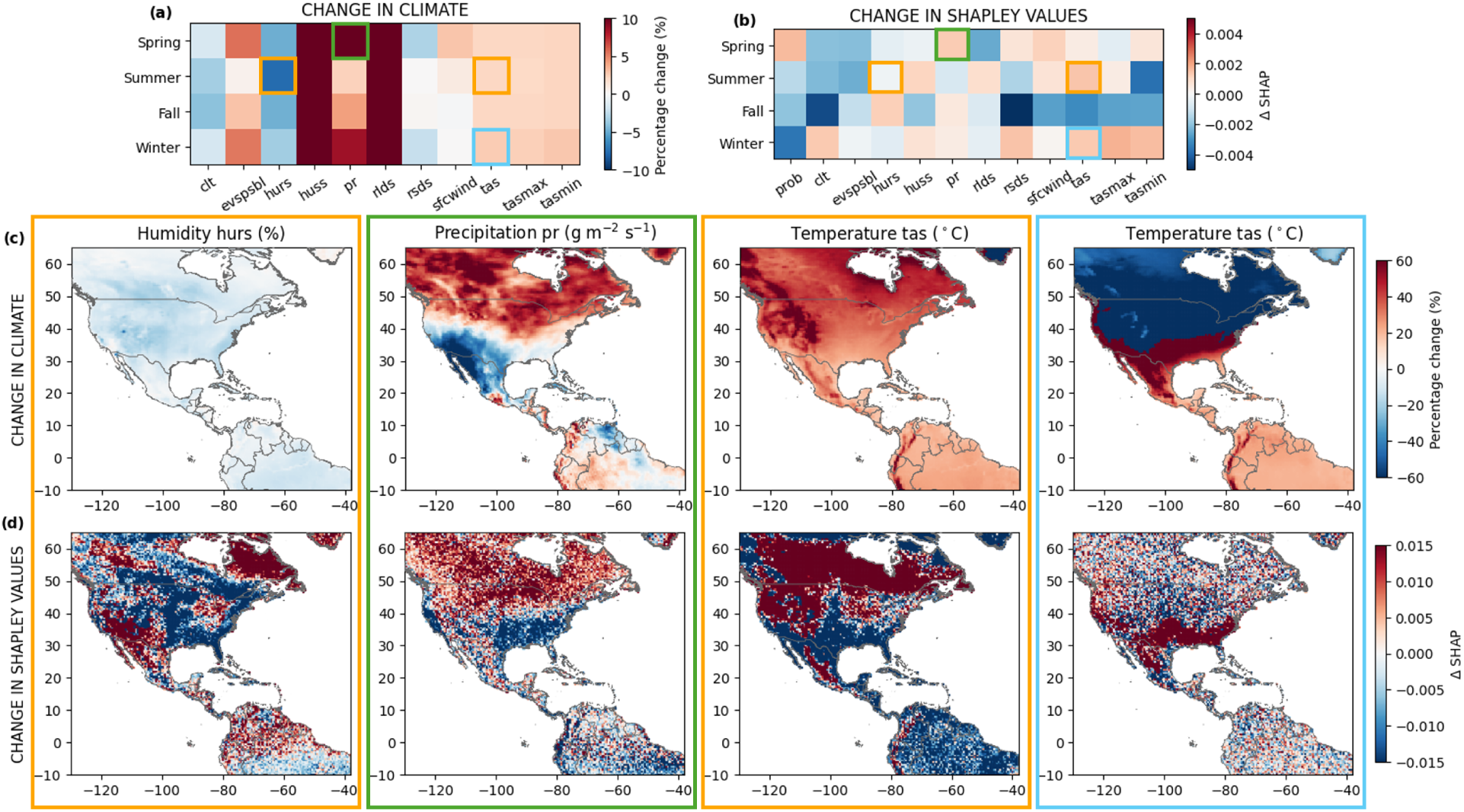
Change in (a) climatic variables and (b) Shapley values between the SSP5-8.5 scenario at the end of the XXI century (2086-2100) and the historical baseline (2010-2024) for each predictor and season. Spatial maps of (c) percentage climatic changes and Δ changes in Shapley values of relative humidity in summer (*hurs*, %), precipitation in spring (*pr*, g m^-2^ s^-1^), and temperature in summer and winter (*tas*, *^◦^*C), compared to 2010-2024. The colored boxes highlight the climatic variables that are investigated. Their color represents the considered seasons: spring (green), summer (orange), fall (red), winter (cyan). Refer to Table A1 for variable abbreviations and full description.

## Discussion

How is climate change likely to alter species ranges and migration patterns? Which environmental variables and climate change scenarios would be responsible for detrimental impacts on the ecology of migratory populations and the key ecosystem services they provide? In this study, we develop a season-aware deep learning approach to model the ecological niche of an iconic migratory species, the monarch butterfly (*Danaus plexippus*), at a pan-American scale. Specifically, our seasonal model can accurately reproduce the full migratory cycle of monarchs across seasons. This framework not only outperforms the conventional time static approach on our held-out test set, but also outperforms it when estimating any season niche from historical data from different climatic conditions compared to those considered in model training. Finally, we predict future shifts of the monarch niche under multiple climate change scenarios. Using Shapley values, an explainable AI method, we establish the contribution of each climatic predictor in shaping *D. plexippus* current and future ecological niche at the global scale in a spatially explicit way.

Our findings suggest that humidity, temperature, and precipitation, in addition to cloud coverage and long-wave radiation, play an important role in shaping the ecological niche of *D. plexippus*. The relevance of these climatic variables for the monarch population size and distribution is in agreement with previous literature [70, 13, 71]. The climatic effects that we highlighted vary in space and in time, with contrasting contributions across the study region and in the different months. Importantly, the contribution of climatic predictors to the winter ecological niche is very distinct from the rest of the year. Given the environmental sensitivity of the wintering diapause state [60, 59], mapping the climatic suitability of overwintering sites is of high interest for butterfly conservation and for long-term sustainability of the host population [66]. At the end of the XXI century, our findings suggest a north-western shift of the monarch niche of [+0.5;+9] degrees in latitude and [-1.5;-6] degrees in longitude, depending on the realized climate change scenario. These distributional changes would be the result of a consistent expansion of suitable areas situated at the northern border of the seasonal ecological niche, counterbalanced by the niche contraction in the southern regions. These losses are expected to involve more than half of the regions currently occupied by resident monarch populations and are mainly located in California and Central America for the summer and winter niche, respectively. The exclusion of these regions from the monarch niche could have disruptive consequences for the population and migratory cycle. Similarly, loss of breeding grounds in the spring-summer season can equally damage monarch population abundance [13, 81]. These considerations regarding predicted range shifts in the monarch niche, in particular for breeding and overwintering grounds, provide essential insights into the monarch migration that should be considered by conservation planning [82].

The study of climate change impacts on species ecological niche faces numerous challenges due to the complex species-specific responses and the simultaneous effect of multiple climatic features. For instance, counteractive climate change impacts on species ranges have already been documented for both plants and animals [83, 84, 85], such as downslope shifts, highlighting the substantial heterogeneity in the climate change effects on species distribution [86]. The study of the ecological niche of migratory species exacerbates the complexity of this issue by adding the temporal component to the niche study [30]. This requires adequate modeling techniques that can disentangle the importance of the multiple climatic effects on species niche in a spatio-temporally dependent manner. The implementation of XAI methods fulfills this need by identifying the relevant predictors linked to the species occurrence in different seasons and responsible for species range shifts [34, 45]. In our work, we use Shapley values to achieve this task [80, 44], and our findings highlight that the same climatic change can have contrasting consequences across the spatial domain and the four seasons. Overall, XAI represents a powerful resource to bridge the gap between ecological relevance for species distribution and powerful black-box modeling tools [87]. Examining changes in Shapley values under different climatic scenarios, we identify whether and which climatic variables would be responsible for the predicted shifts, both globally and locally, for the monarch butterfly niche.

Although bringing novel insights into the biogeography of migratory species under climate change, our study is not exempt from limitations. Statistical modeling frameworks for species distribution have a limited predictive capability in very different climatic settings [88]. Although we test that our approach works well for past climatic conditions, this does not exclude inaccuracies when the most extreme climatic scenarios are examined. Importantly, our considerations are based on the monarch response to current and past environmental conditions, lacking the inclusion of potential evolutionary dynamics that would drive adaptation to future climate. In addition, true species absences should be used in model training and validation, although those types of data are limited compared to presence-only observations from which we derived pseudoabsences [89]. While we explicitly incorporate the intra-annual (seasonal) variability of climatic variables and the consequentiality of seasonal niches, the climatic inter-annual differences within the considered temporal intervals are neglected. A possible way to integrate yearly dynamics could be to implement our seasonal model architecture year by year and to obtain a more precise niche estimate over time. However, this annual analysis would be more suited to short-term forecasting rather than long-term estimates of climate change impacts.

Our modeling approach advances current knowledge on the species’ climatic suitability, but needs to be paired with a population ecological model to investigate the effective impacts of climate change on monarch population abundance and survival. In addition, our framework considers the effect of multiple climatic features on the distribution of monarch butterflies, disregarding biotic interspecific ecological interactions. *D. plexippus* larvae feed on specific host plants, i.e. milkweeds (*Asclepias* spp.) [90], whose population is experiencing a widespread decline due to the intensive use of glyphosate herbicides for agriculture [13]. The decreasing abundance of native milkweed threatens the monarch population to be captured in a trophic ecological trap: the non-native and invasive milkweed (*Asclepias curassavica*) is increasing the sedentary behavior of the monarch population by providing a year-round food source, but of lower quality that can negatively impact butterfly mass and survival [91]. Therefore, monarch ecological niche shifts under global warming should be evaluated in relation to milkweeds, natural enemies [92] and infectious diseases, such as the devastating protozoa *Ophryocystis elektroscirrha*. Deep learning approaches like ours represent a flexible and valid option to embed these multi-trophic interactions and their potential effects on monarch distribution. We leave these studies for future work.

Although the monarch butterfly presents a unique migration pattern and climatic dependencies specific to its population ecology, it is not the only species that needs to travel through a warming world: migration is a widespread behavior in the animal kingdom, with more than 600 migratory species only for butterflies [93]. Therefore, our approach has the potential to advance scientific understanding regarding the ecological niche of other migratory animals and their vulnerability to global warming [10]. Investigating the current and future challenges that climate change poses to species macroecology and biogeography is essential to implement effective conservation strategies aimed at preserving biodiversity and ecosystem health.

## Appendix

**Table A1:**
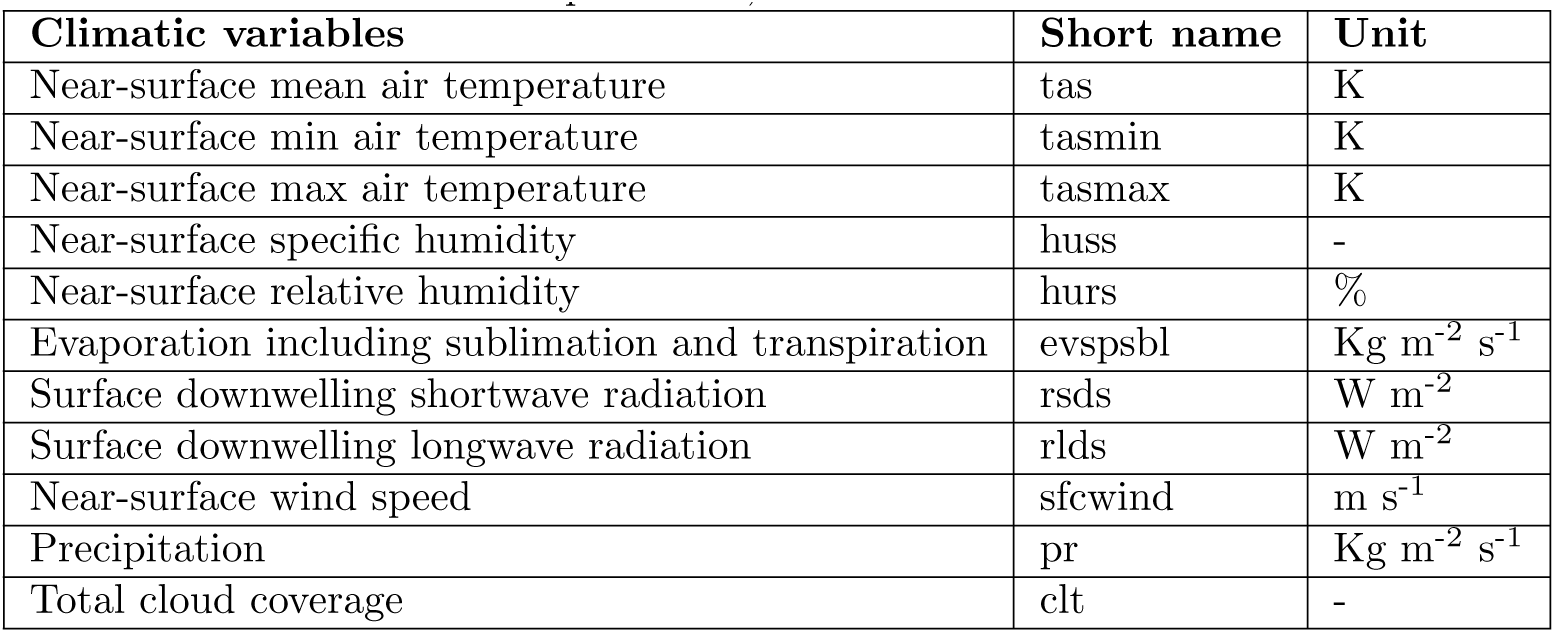
Climatic predictors, short names and unit of measure.

**Table A2:** Global (any season) and seasonal MaxEnt (linear, hinge and product feature classes) model performance with both of the Area Under the ROC Curve (AUC_ROC_), the Area Under the Precision-Recall Curve (AUC_PR_), and the True Skill Statistic (TSS) using the test dataset of 2010-2024. M0 performance is reported for comparison.

| Model | Performance | $AUC_{ROC}$ | $AUC_{PR}$ | TSS |
| --- | --- | --- | --- | --- |
| MaxEnt | Global | 0.878 | 0.920 | 0.666 |
|  | Spring | 0.874 | 0.910 | 0.683 |
|  | Summer | 0.891 | 0.979 | 0.678 |
|  | Fall | 0.889 | 0.971 | 0.672 |
|  | Winter | 0.860 | 0.823 | 0.631 |
| M0 | Global | 0.874 | 0.903 | 0.683 |
|  | Spring | 0.862 | 0.893 | 0.688 |
|  | Summer | 0.907 | 0.980 | 0.687 |
|  | Fall | 0.886 | 0.966 | 0.684 |
|  | Winter | 0.840 | 0.771 | 0.674 |

**Figure A1:**
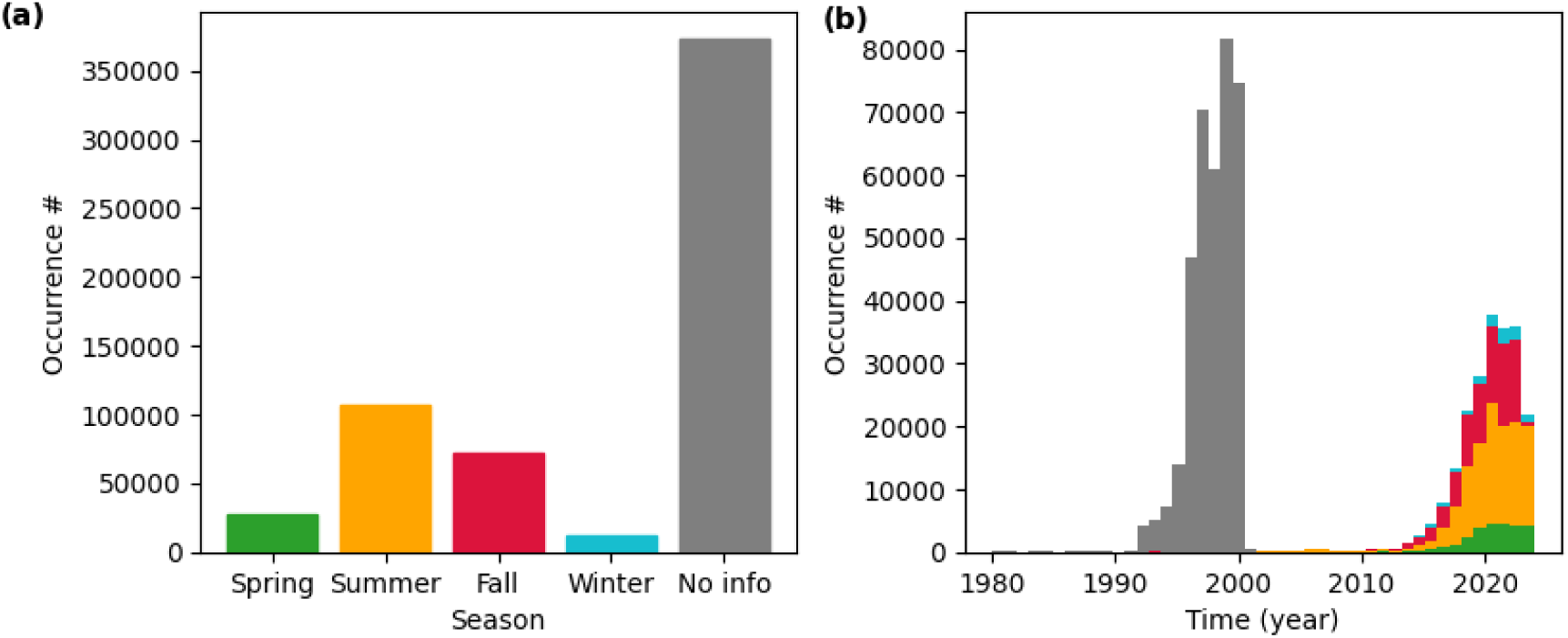
Monarch butterfly occurrence data: (a) total number of observations and (b) distribution in time of available observations in spring (green), summer (orange), fall (red), winter (cyan), and for samples without seasonal information (gray).

**Figure A2:**
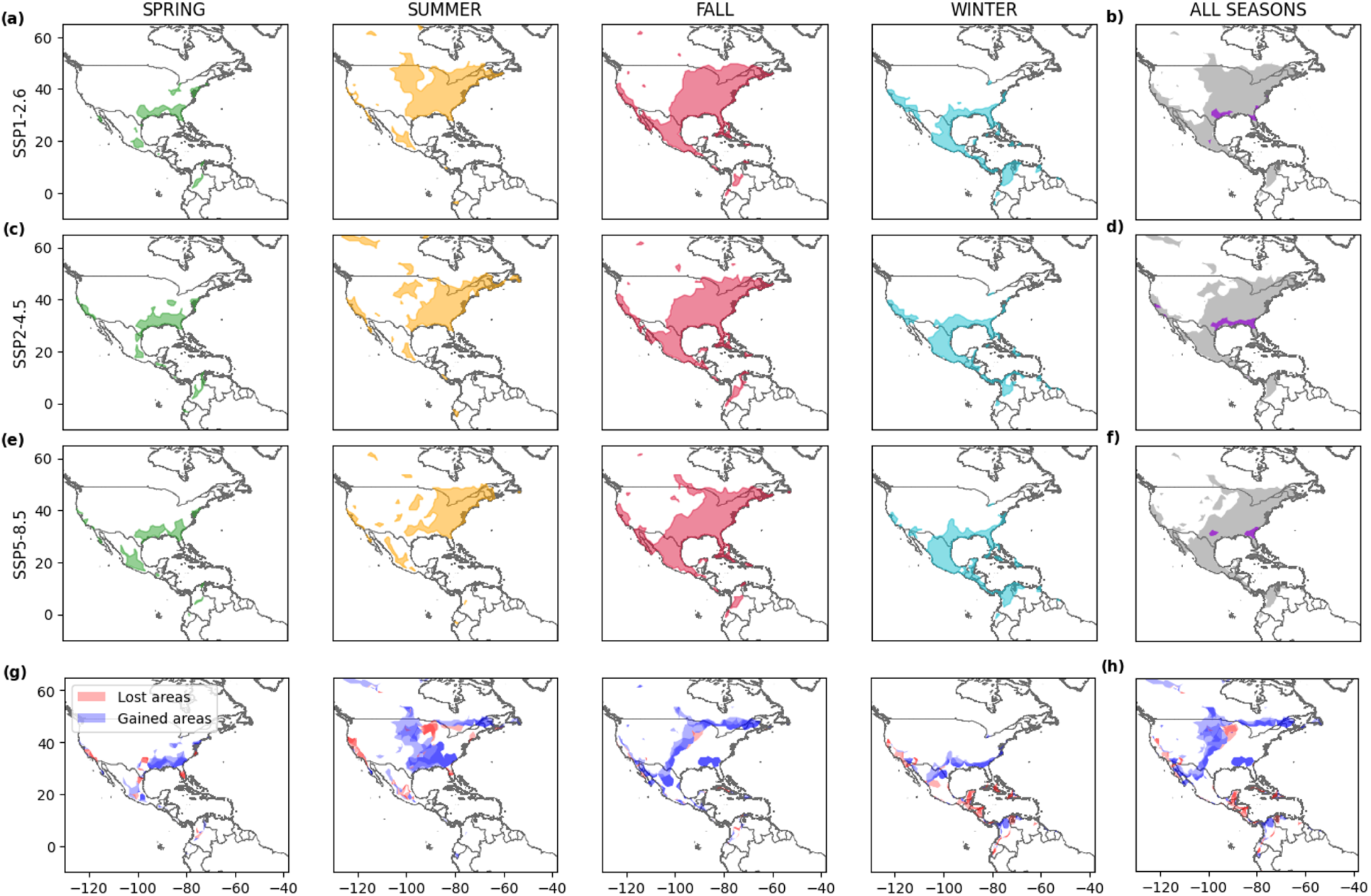
Projected seasonal seasonal (a, c, e) and global (all seasons, b, d, f) niches of the monarch butterfly (spring in green, summer in orange, fall in red, winter in cyan, any season in gray and year-round in purple) at the middle of the XXI century (**2040-2054**) under SSP1-2.6 (a, b), SSP2-4.5 (c, d) and SSP5-8.5 (e, f) climate change scenarios. Lost (red) and gained (blue) areas, compared to 2010-2024, are represented for each season (g) and for all seasons (h). Darker areas (panels g, h) represent regions of agreement between the three scenarios.

**Figure A3:**
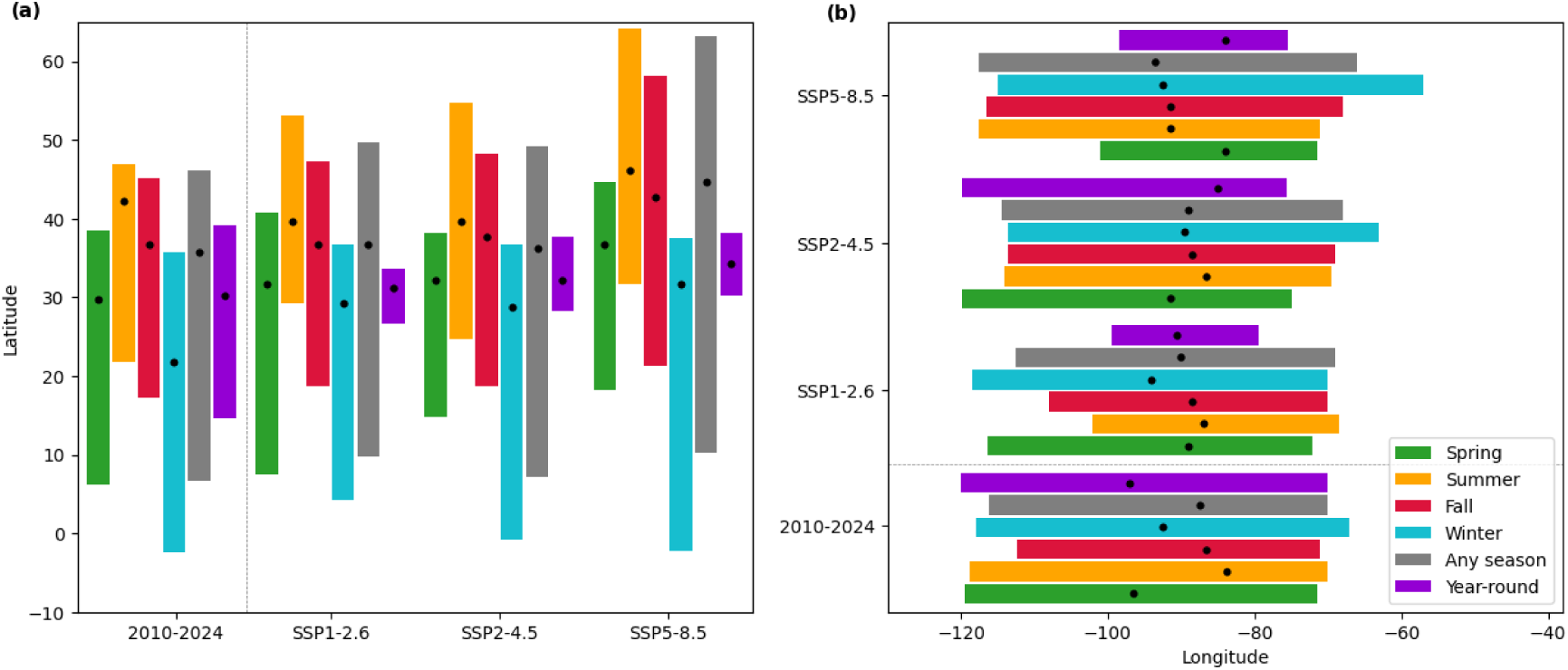
Latitudinal (a) and longitudinal (b) distribution of the monarch butterfly niche (spring in green, summer in orange, fall in red, winter in cyan, any season in gray and year-round in purple) in 2010-2024 and at the end of the XXI century (2086-2100) under SSP1-2.6, SSP2-4.5 and SSP5-8.5 climate change scenarios. The bars are plotted within the 5^th^ and the 95^th^ percentile of latitude and longitude, while the black dots represent the medians.

**Figure A4:**
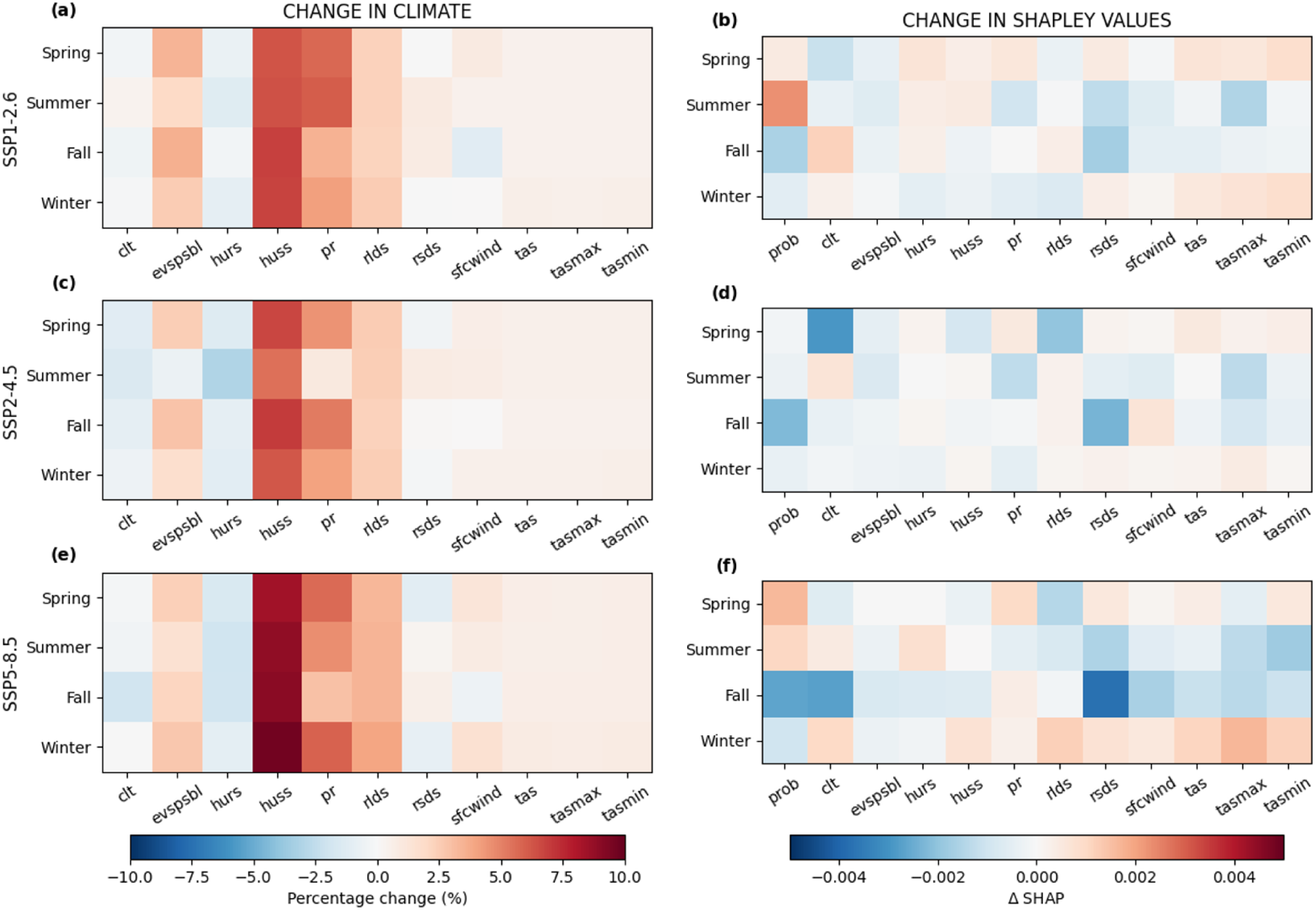
Change in (a, c, e) climatic variables and (b, d, f) Shapley values between the SSP1-2.6 (a, b), SSP2-4.5 (c, d), and SSP5-8.5 (e, f) scenario at the middle of the XXI century (**2040-2054**) and the historical baseline (2010-2024) for each predictor and season.

**Figure A5:**
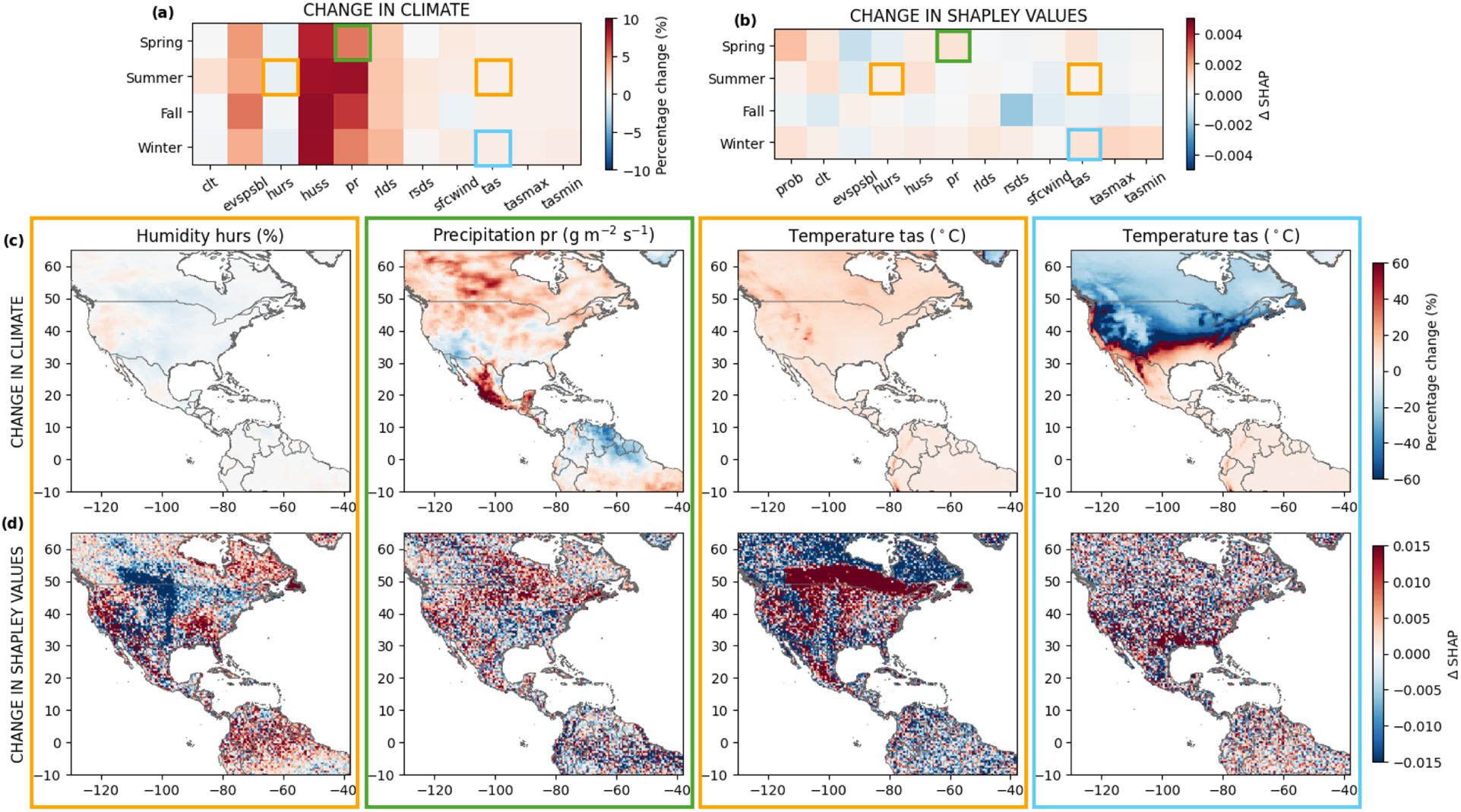
Change in (a) climatic variables and (b) Shapley values between the **SSP1-2.6** scenario at the end of the XXI century (2086-2100) and the historical baseline (2010-2024) for each predictor and season. Spatial maps of (c) percentage climatic changes and Δ changes in Shapley values of relative humidity in summer (*hurs*, %), precipitation in spring (*pr*, g m^-2^ s^-1^), and temperature in summer and winter (*tas*, *^◦^*C), compared to 2010-2024. The colored boxes highlight the climatic variables that are investigated. Their color represents the considered seasons: spring (green), summer (orange), fall (red), winter (cyan). Refer to Table A1 for variable abbreviations and full description.

**Figure A6:**
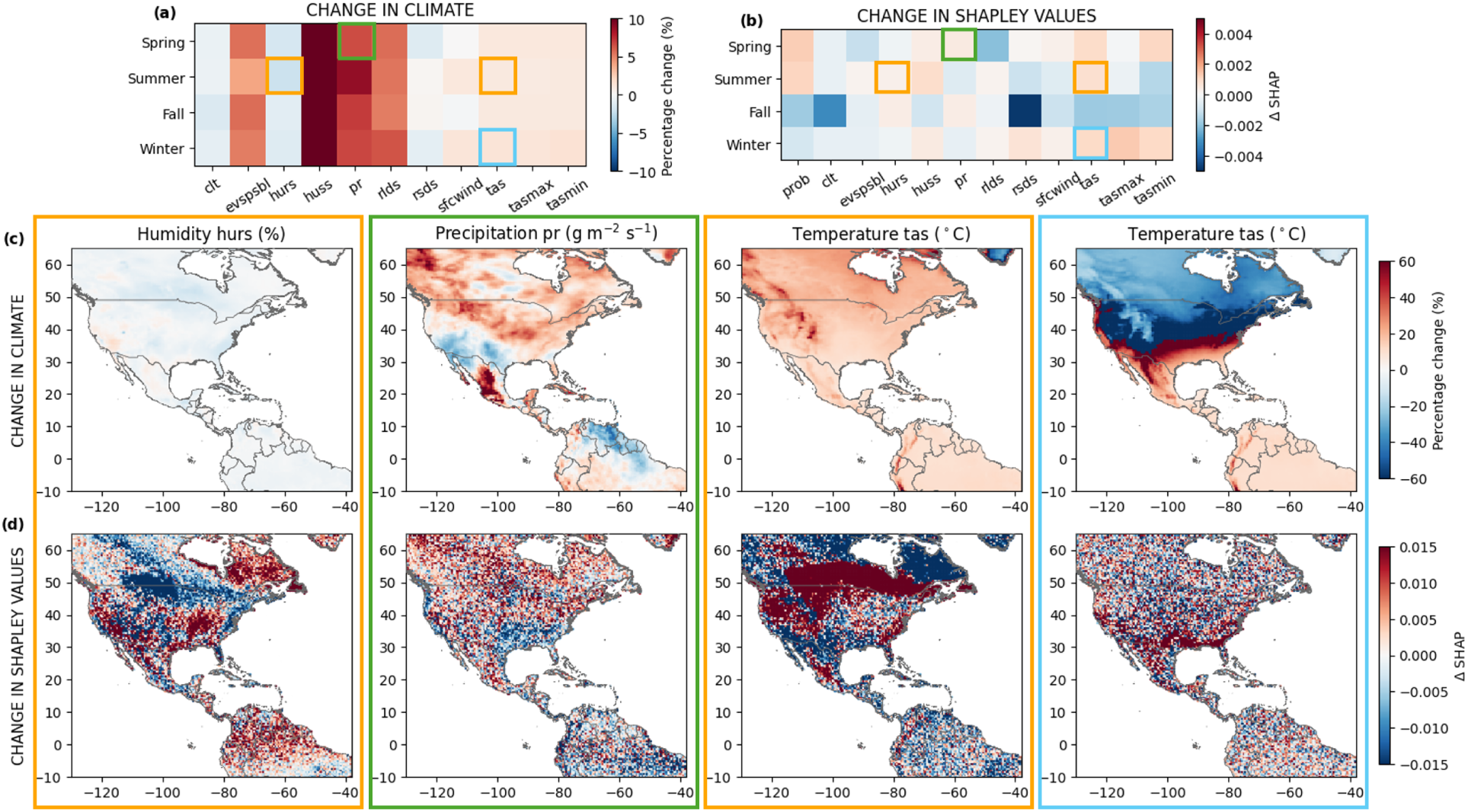
Change in (a) climatic variables and (b) Shapley values between the **SSP2-4.5** scenario at the end of the XXI century (2086-2100) and the historical baseline (2010-2024) for each predictor and season. Spatial maps of (c) percentage climatic changes and Δ changes in Shapley values of relative humidity in summer (*hurs*, %), precipitation in spring (*pr*, g m^-2^ s^-1^), and temperature in summer and winter (*tas*, *^◦^*C), compared to 2010-2024. The colored boxes highlight the climatic variables that are investigated. Their color represents the considered seasons: spring (green), summer (orange), fall (red), winter (cyan). Refer to Table A1 for variable abbreviations and full description.

## Notes

### Competing Interest Statement

The authors have declared no competing interest.

### Funding

- Swiss National Science Foundation: 204057

